# Development and applications of a CRISPR activation system for facile genetic overexpression in *Candida albicans*

**DOI:** 10.1101/2022.08.15.501889

**Authors:** Nicholas C. Gervais, Alyssa A. La Bella, Lauren F. Wensing, Jehoshua Sharma, Victoria Acquaviva, Madison Best, Ricardo Omar Cadena López, Meea Fogal, Deeva Uthayakumar, Alejandro Chavez, Felipe Santiago-Tirado, Ana L. Flores-Mireles, Rebecca S. Shapiro

## Abstract

For the fungal pathogen *Candida albicans*, genetic overexpression readily occurs via a diversity of genomic alterations, such as aneuploidy and gain-of-function mutations, with important consequences for host adaptation, virulence, and evolution of antifungal drug resistance. Given the important role of overexpression on *C. albicans* biology, it is critical to develop and harness tools that enable the analysis of genes expressed at high levels in the fungal cell. Here, we describe the development, optimization, and application of a novel, single-plasmid-based CRISPR activation (CRISPRa) platform for targeted genetic overexpression in *C. albicans*, which employs a guide RNA to target an activator complex to the promoter region of a gene of interest, thus driving transcriptional expression of that gene. Using this system, we demonstrate the ability of CRISPRa to drive high levels of gene expression in *C. albicans*, and we assess optimal guide RNA targeting for robust and constitutive overexpression. We further demonstrate the specificity of the system via RNA sequencing. We highlight the application of CRISPRa to overexpress genes involved in pathogenesis and drug resistance and contribute towards the identification of novel phenotypes. Together, this tool will facilitate a broad range of applications for the study of *C. albicans* genetic overexpression.

## Introduction

Fungal pathogens are an increasingly important cause of human illness, with significant impacts on morbidity and mortality (1). *Candida albicans* is an opportunistic fungal pathogen that inhabits the microbiome of most healthy adults as a commensal organism (2, 3). However, it is also responsible for invasive infections, particularly amongst immunocompromised individuals, which are often fatal (4). Instances of *C. albicans* infections are becoming more prevalent globally, in part due to an increasing population of susceptible elderly and immunocompromised individuals (4). Illustratively, systemic *Candida* infections have been identified in up to 15% of patients hospitalized for COVID-19, and patients with invasive candidiasis spent longer in the hospital Intensive Care Unit (ICU) on average, and had more severe symptoms than those COVID-19 patients without a *Candida* infection (5). The economic cost of *Candida* infections is estimated to be ~1.4 billion dollars yearly in the United States alone, hospitalizing more than 26,000 people (6). With limited development of novel antifungal drugs (7) and increasing prevalence of antifungal drug-resistant strains and species (8–10), *Candida* pathogens remain a serious threat to human health.

Given the importance of *C. albicans* and other fungal pathogens, it is imperative that tools are developed and applied to manipulate these organisms and dissect genetic functions. For *C. albicans*, along with countless other microbial species, CRISPR-based genetic manipulation tools have revolutionized the ability of scientists to modify and study gene function (11, 12). Such CRISPR platforms typically rely on a CRISPR-associated endonuclease protein, often Cas9, targeted to a genomic region of interest via a single guide RNA (sgRNA), resulting in targeted DNA double-strand breaks (DSBs), and subsequent repair via non-homologous end-joining or homology-directed repair via a donor DNA repair template (13, 14). Since the initial development of CRISPR tools for genetic editing in *C. albicans* (15), numerous systems have been developed to modify this fungal genome with efficiency and specificity (11, 16–24), and to build and screen genetic mutant libraries (24, 25). Together, these gene editing systems have played an important role in providing novel biological insight into this fungal pathogen.

In addition to these canonical CRISPR-based genetic modification systems that introduce mutations, insertions, or deletions into the genome, alternative CRISPR technologies have been developed that modulate the expression of *C. albicans* genes (26, 27). These CRISPR systems rely on the inactivation of the Cas9 endonuclease, to generate a nuclease-dead (dCas9) protein, which can be fused to constructs from a transcriptional repressor or activator that generate CRISPR interference (CRISPRi) or CRISPR activation (CRISPRa) systems, respectively (28, 29). These dCas9 fusions, when targeted to the promoter region of a gene via the sgRNA, can repress or activate gene transcription from a gene of interest, and have been widely applied in diverse organisms (14, 30–34). In *C. albicans*, these modified CRISPR systems have enabled the modulation and study of important fungal genes, including essential genes (26, 27, 35).

While a majority of CRISPR systems and other genetic manipulation platforms focus on gene mutation, deletion, or downregulation, it is also critical to develop techniques that enable the upregulation or overexpression of genes. Indeed, genetic overexpression in *C. albicans* occurs readily via gene copy number amplification and aneuploidy, and these copy number amplification events are frequently observed in genetically diverse clinical isolates (36–38). The upregulation of gene expression plays a critical role in diverse aspects of fungal biology (39), including the evolution of antifungal drug resistance (40–45), host colonization (46, 47), and virulence (41, 48). Additionally, in other microbial systems, including bacteria, protozoa, and the model yeast *Saccharomyces cerevisiae*, genetic overexpression libraries have proven to be critical tools for identifying targets of drugs with uncharacterized mechanisms of action, as mutants overexpressing the drug target are typically more resistant to the drug in question (49–52). Overexpression screens can also help dissect molecular pathways (53–55). In *C. albicans*, several techniques exist for genetic overexpression (39), including promoter replacement strategies (55–58), the generation of ORFeome collection strains (59, 60), and CRISPR-based methods (27). While these these tools have made important contributions to understanding overexpression in *C. albicans*, there remains a need for simple and rapid methods to efficiently target fungal genes of interest for overexpression, and to do so in diverse strain backgrounds.

Here, we introduce a CRISPR-based tool for genetic overexpression in *C. albicans*, to bolster the existing functional genetic toolbox available to manipulate genes in this critical fungal pathogen. Our CRISPRa tool exploits a *C. albicans*-optimized tripartite activator complex fused to dCas9, which was previously developed for highly-efficient transcriptional regulation in *S. cerevisiae* (61). This CRISPRa system is distinct from other CRISPR-based gene activation systems for *C. albicans*, as it is an efficient single-plasmid system designed for rapid Golden Gate cloning and facile fungal strain generation, which can be readily applied to diverse strain backgrounds including clinical isolates. We test CRISPRa guide targeting principles based on predicted transcriptional start sites, confirm on-target guide efficiency, and demonstrate the ability of this CRISPRa system to drive high levels of expression of *C. albicans* genes. Using this optimized CRISPRa technique, we validate its ability to drive overexpression of genes with established roles in antifungal drug resistance and biofilm formation. Together, this work introduces a novel CRISPRa system for genetic overexpression in *C. albicans*, with a wide range of future applications.

## Results

### Development of a single-plasmid CRISPRa tool in C. albicans

First, we designed a single-plasmid CRISPRa system for targeted gene activation, optimized for *C. albicans*. We exploited an integrating plasmid backbone, which we previously used for CRISPRi in *C. albicans*, containing a codon-optimized dCas9 and Golden Gate assembly-compatible sgRNA cloning site (26). This plasmid can be linearized to integrate at the *C. albicans NEUT5L* locus (a large intergenic region whose disruption does not affect fungal fitness (62)), and contains a dual SapI cloning site for facile and highly-efficient Golden Gate cloning of the sgRNA targeting sequence. This plasmid also contains a dominant nourseothricin resistance (NATr) cassette, to enable strain generation in a diversity of fungal strains, including clinical isolates.

To generate a version of dCas9 that could lead to strong constitutive activation of genes of interest, we generated a plasmid with dCas9 fused to the tripartite activator complex, VP64-p65-Rta (VPR) (**FIGURE 1a,b**). VPR consists of three previously-described transcriptional activators: (1) VP64, a viral transcriptional activator domain composed of four tandem copies of VP16 (Herpes Simplex Viral Protein 16), which has been used extensively in CRISPRa systems (63); (2) p65, the NF-κB subunit responsible for the strong transcription activation in mammalian cells (64); and (3) Rta, the Epstein-Barr virus R transactivator, with strong transcriptional activation properties (65) (**FIGURE 1a, b**). Together the VPR activator complex fused to dCas9 and targetted to the promoter region of genes has previously been demonstrated to generate strong gene activation in *S. cerevisiae* (61). Exploiting the ability of dCas9 to bind and be targeted to genomic loci via sgRNAs enables us to design sgRNAs that will target dCas9 along with these strong transcriptional activators to any gene promoter of interest.

**Figure 1.**
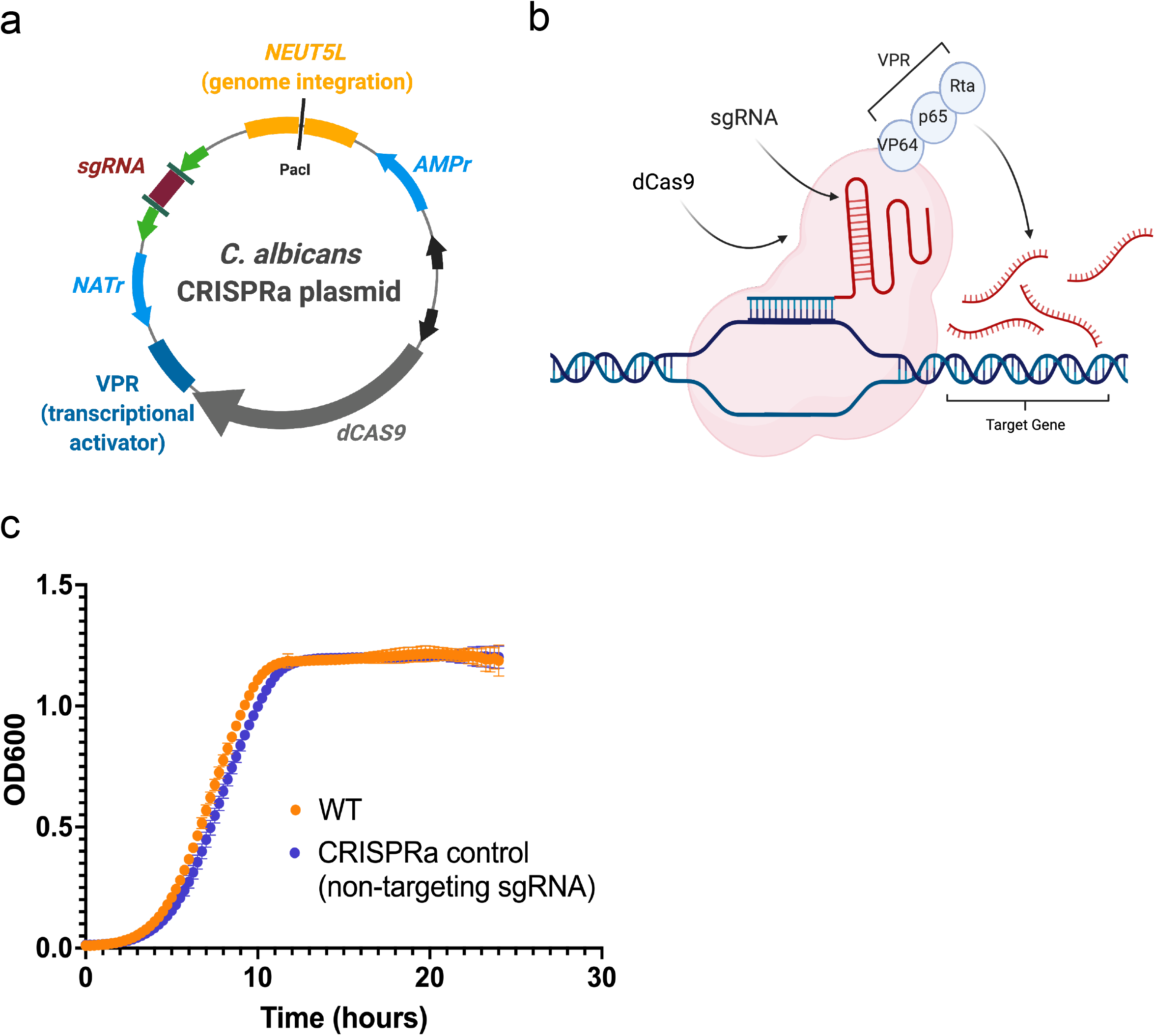
Design of a CRISPRa system for *C. albicans*. **(a)**The single plasmid system contains a catalytically inactive (“dead”) dCas9 protein, paired with the VPR (VP64-P65-Rta) complex of activator domains to cause upregulation at a target locus. The single-guide RNA (sgRNA) region can be subjected to Golden Gate cloning with a custom N20 sequence in order to target the dCas9-VPR complex to a desired region of DNA through complementary base pairing. The PacI enzyme is used to linearize the plasmid at the *NEUT5L* locus, which allows the plasmid to be integrated into the cell’s chromosome and remain in the cell following cell division. Panel created with BioRender.com. **(b)** Once the sgRNA recognizes and binds to a target sequence of DNA, the dCas9-VPR complex will cause the recruitment of transcription initiation complex factors to the adjacent genetic material, thus increasing gene expression from that region. Created with BioRender.com. **(c)** Wild-type *C. albicans* cells and cells containing a non-targeting CRISPRa plasmid were grown in YPD at 30°C to compare fitness under standard laboratory conditions. Growth was recorded via optical density at 600nm at 15 min intervals over the course of 24h. Data points show the mean and standard deviation at each time point (n = 32).

This plasmid was generated by homology-based cloning and validated via Sanger sequencing, and has been deposited to Addgene (**TABLE S1**, Addgene plasmid #182707). To confirm if the expression of the dCas9-VPR construct had an impact on fungal fitness, we compared the growth of a wild-type *C. albicans* strain with a strain transformed with the dCas9-VPR CRISPRa plasmid (expressing a non-targeting sgRNA: a random sequence of nucleotides that have no predicted complementarity to any region in the *C. albicans* genome). We found no difference in growth between these strains (**FIGURE 1c**), suggesting no significant impact of the CRISPRa system on *C. albicans* fitness.

### Validation and sgRNA design principles for CRISPRa overexpression in C. albicans

Next, we aimed to validate that the CRISPRa system could successfully overexpress genes in *C. albicans*, and sought to determine the optimal sgRNA design strategy to maximize genetic regulation. Our previous *C. albicans* CRISPRi system exploited sgRNAs ~ 50 - 150bp upstream of the start codon to functionally repress genes of interest (26). However, CRISPRa systems are known to have different optimal sgRNA targeting regions (66, 67), and our previous design did not take into account the transcription start site (TSS), which is known to be an important predictor of regulatory efficiency for CRISPRi/a systems (66, 68), but which had not been robustly characterized across all *C. albicans* genes at the time. More recently, the TSS for many *C. albicans* genes has been predicted (69). Thus, we sought to determine the optimal design for CRISPRa sgRNAs based on the start codon and the TSS. We identified three *C. albicans* genes with a predicted TSS that was particularly far upstream of the start codon compared to all other genes in the genome (801-952 bp upstream): *KSP1, ALS3*, and *FGR28*. We designed six sgRNAs for each gene, targeting upstream of the start codon (as previously designed for CRISPRi (26)), as well as upstream of the TSS (based on predicted TSS (69)).

We generated CRISPRa plasmids based on these sgRNAs, used these to create *C. albicans* strains, and monitored overexpression of each gene via quantitative reverse-transcription PCR (RT-qPCR). This analysis validated our novel CRISPRa system and its ability to overexpress *C. albicans* genes ~ 2 - 12-fold, based on different sgRNA designs and target genes (**FIGURE 2**). We found that in some instances, such as for *KSP1*, the TSS was in fact a better predictor of CRISPRa overexpression, as the most robust overexpression of this gene was observed with sgRNAs targetted upstream of the TSS, while no overexpression was observed based on targeting upstream of the start codon (**FIGURE 2a**). In other instances, such as for *ALS3*, sgRNA targeting upstream of the start codon led to very high levels of overexpression, while sgRNAs targeted to the TSS region led to moderate overexpression (**FIGURE 2b**). For *FGR28*, modest overexpression (~2-fold) was achieved with sgRNAs targeted to either the start codon or the TSS upstream regions (**FIGURE 2c**), together suggesting that optimal sgRNA design can be predicted based on targeting the TSS and/or start codon, but varies by gene. This data also suggests that our CRISPRa tool may have additional applications in the functional assessment of key transcriptional regulatory regions in *C. albicans*. These differences in sgRNA targeting may be based on differences in overall sgRNA efficiency or may be attributed to inaccurate TSS predictions, or complex promoter dynamics and the preferential use of different TSS regions in different conditions (70).

**Figure 2.**
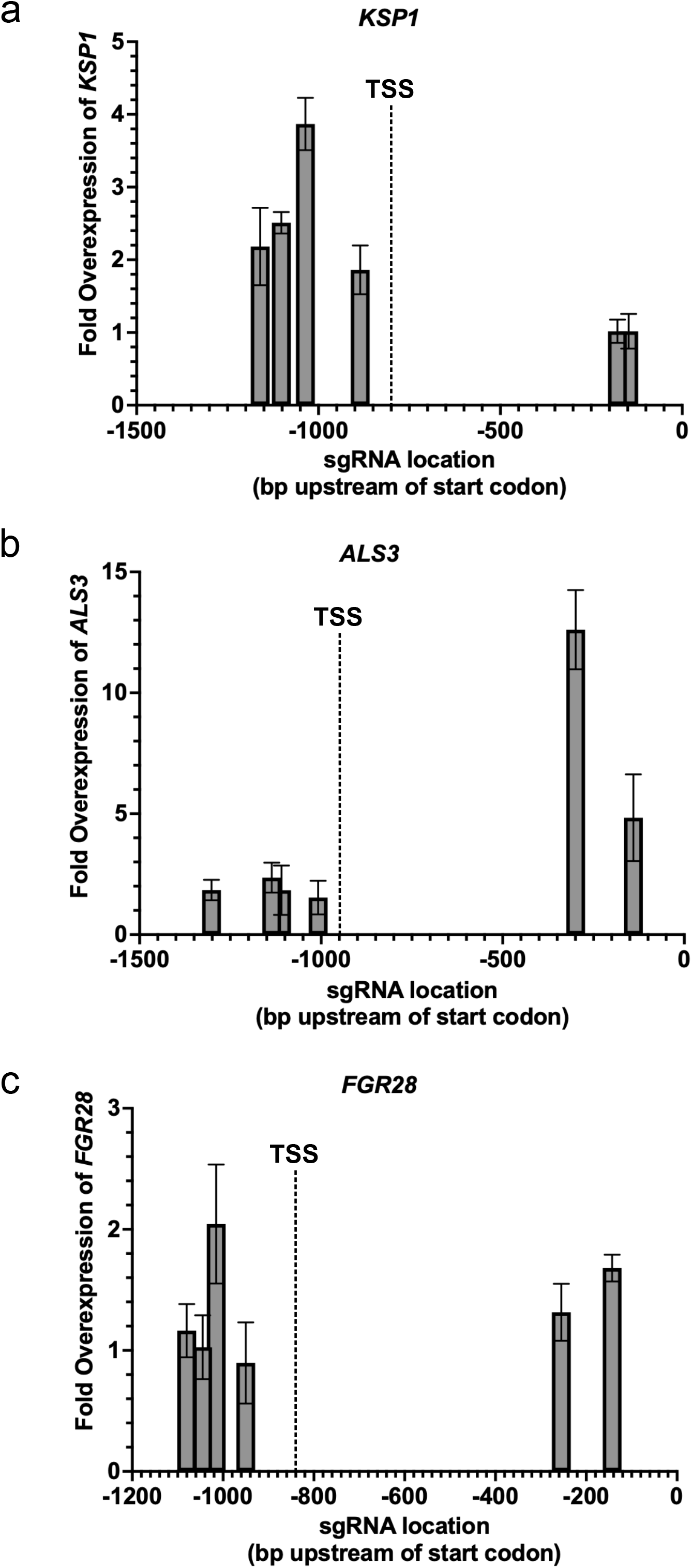
Determining optimal guide-design principles for CRISPRa overexpression in *C. albicans*. CRISPRa strains were created using six different sgRNA molecules designed to target different positions in the promoter region of each of **(a)** *KSP1*, **(b)** *ALS3*, and **(c)** *FGR28* upstream of either the start codon or the predicted dominant transcriptional start site (TSS). Fold-change in the expression of the target gene relative to the housekeeping gene *ACT1* in the experimental strains and the non-targeting CRISPRa control strain was measured with RT-qPCR and compared. Data points represent the mean fold-difference and standard deviation in expression of the respective target gene in the CRISPRa strains compared to the non-targeting CRISPRa control (n = 3).

### Use of CRISPRa to study antifungal drug resistance phenotypes

To validate that the CRISPRa system could recapitulate phenotypes associated with *C. albicans* gene overexpression, we targeted genes with known phenotypes associated with high levels of expression. Initially, we focused on *CDR1*, a well-characterized *C. albicans* gene, which encodes a well-characterized drug efflux pump (71), and whose overexpression is known to play a role in drug resistance to azole antifungals in laboratory and clinical isolates (72–78). To determine an optimal range in which to target our sgRNA in the *CDR1* promoter, we designed three sgRNAs targeting loci approximately 90 bp, 200 bp, and 290 bp upstream of the *CDR1* TSS. These sgRNAs were each cloned into our CRISPRa dCas9-VPR backbone and transformed into *C. albicans*. Next, we quantified the expression of *CDR1* in the three different fungal strains via RT-qPCR, compared with a control strain containing the CRISPRa plasmid with a non-targeting sgRNA. We found that targeting the dCas9-VPR construct to the *CDR1* promoter enhanced transcription by 2.9 - 5.6 fold (**FIGURE 3a**), indicating that this CRISPRa system is indeed able to drive enhanced levels of gene expression from a target gene of interest. For *CDR1*, we found the highest level of overexpression was obtained by targeting the dCas9-VPR construct ~200 bp upstream of the TSS (**FIGURE 3a**), which helped inform the design of sgRNAs for subsequent target genes.

**Figure 3.**
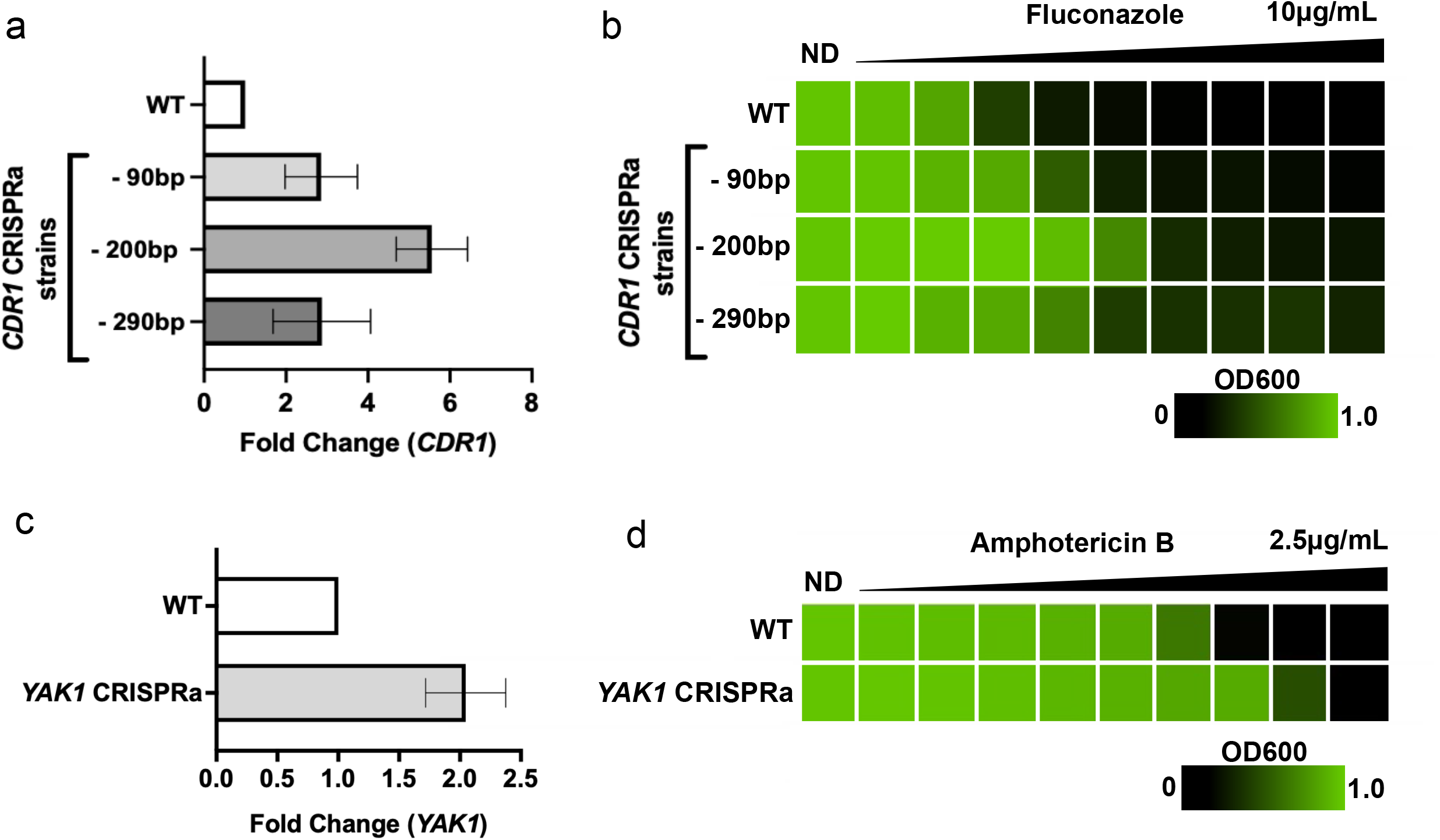
Using CRISPRa to validate antifungal drug resistance phenotypes. **(a)** Three *C. albicans* CRISPRa strains were created targeting the drug transporter gene *CDR1* for overexpression. Fold-change in the expression of *CDR1* relative to the housekeeping gene *ACT1* in the three experimental strains and the non-targeting CRISPRa control strain was measured with RT-qPCR. All three experimental strains are represented by the position of their sgRNA in the promoter region upstream of the proposed dominant *CDR1* transcriptional start site. Data represent the mean fold-difference and standard deviation in expression of *CDR1* in the CRISPRa strains compared to the non-targeting CRISPRa control (n = 3). **(b)** The fitness of all three CRISPRa strains targeting *CDR1* for overexpression was measured in the presence of the antifungal drug fluconazole with a minimum inhibitory concentration assay. The OD600 value of each strain in each concentration of fluconazole was measured to represent growth in differing conditions, where OD600 values approaching 1.0 reflect strong fitness. **(c)** Fold-change in the expression of *YAK1* relative to the housekeeping gene *ACT1* in the experimental strain and the non-targeting CRISPRa control strain was measured with RT-qPCR. Data represent the mean fold-difference and standard deviation in expression of *YAK1* in the CRISPRa strains compared to the non-targeting CRISPRa control (n = 3). **(d)** The fitness of the CRISPRa strain targeting *YAK1* for overexpression was measured in the presence of the antifungal drug amphotericin B with a minimum inhibitory concentration assay. The OD600 value of each strain in each concentration of amphotericin B was measured to represent growth in differing conditions, where OD600 values approaching 1.0 reflect strong fitness.

In order to determine if *CDR1*-overexpressing strains had measurable changes in antifungal drug resistance phenotypes, we performed minimum inhibitory concentration (MIC) assays with these strains in the presence of the antifungal drug fluconazole. We found that strains overexpressing *CDR1* exhibited increased resistance to fluconazole compared with the control strain and that higher levels of overexpression corresponded with higher levels of resistance (**FIGURE 3b**). Together, this confirms that our CRISPRa system can be exploited to functionally overexpress target genes of interest, and recapitulate phenotypes associated with high levels of gene expression. Further, targeting dCas9-VPR to different promoter loci leads to variable levels of overexpression, which could be used to study correlations between expression levels and associated phenotypes.

We next sought to determine the ability of our CRISPRa system to be employed to characterize a mutant strain that, to our knowledge, has never been profiled with regard to antifungal drug susceptibility. We selected the gene *YAK1* as a target for overexpression with CRISPRa. *C. albicans YAK1* is a predicted serine-threonine protein kinase, inactivation of which renders cells more sensitive to the antifungal amphotericin B (79). While *YAK1* overexpression has been shown to promote filamentation in *C. albicans* (80), and there is emerging evidence for a direct relationship between the two phenotypes (81), the role of overexpressing this factor in antifungal drug resistance has not previously been established. We generated a CRISPRa strain targeting *YAK1* for overexpression (**FIGURE 3c**) and profiled the ability of this strain to grow in the presence of amphotericin B. We found that the *YAK1* CRISPRa strain was more resistant to amphotericin B based on MIC testing on this strain compared to the control strain (**FIGURE 3d**). This highlights our capacity to exploit this CRISPRa system to investigate novel phenotypes in *C. albicans*.

### Validation of CRISPRa overexpression targeting fungal pathogenesis traits

Next, we aimed to validate the CRISPRa system as a means to assess *C. albicans* genes involved in fungal pathogenesis traits. We focused on *ALS1*, a well-characterized *C. albicans* adhesin gene, with important roles in adherence, biofilm formation, and virulence (25, 82–86). Previous work has demonstrated that overexpression of *ALS1* via promoter replacement increases fungal adherence (83), therefore we chose to overexpress this gene using our CRISPRa system. We designed and cloned two sgRNAs targeting approximately 130bp and 260bp upstream of the *ALS1* TSS, and generated *C. albicans ALS1* CRISPRa strains with each of these constructs. We performed RT-qPCR to confirm the overexpression of these strains, which enhanced the expression of *ALS1* by ~25 and ~6 fold, respectively (**FIGURE 4a**).

**Figure 4.**
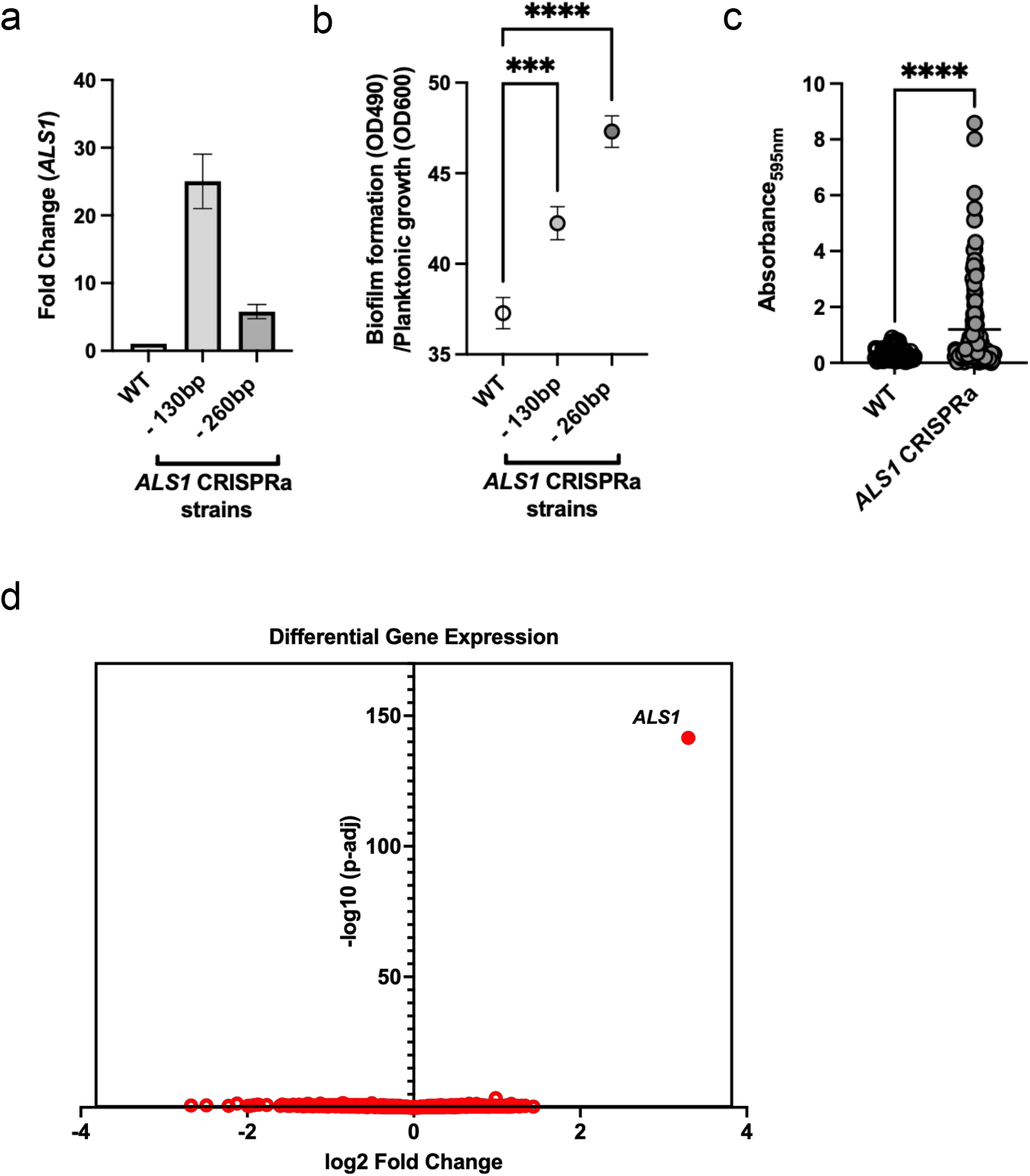
Validation of CRISPRa overexpression and target-specificity through targeting fungal pathogenesis traits. **(a)** Two *C. albicans* CRISPRa strains targeting different positions in the promoter region of the adhesin protein gene *ALS1* were created. Fold-change in the expression of *ALS1* relative to the housekeeping gene *ACT1* in both experimental strains and the non-targeting CRISPRa control strain was measured with RT-qPCR. Experimental strains are represented by the position of their sgRNA in the promoter region upstream of the proposed dominant *ALS1* transcriptional start site. Data points represent the mean fold-difference and standard deviation in expression of *ALS1* in the CRISPRa strains compared to the non-targeting CRISPRa control (n = 3). **(b)** Two CRISPRa strains targeting *ALS1* and a non-targeting CRISPRa control strain were grown in the biofilm-inducing media RPMI and left to grow statically at 37°C for 24h to allow biofilms to form. The robustness of biofilms formed by all strains was elucidated through an XTT reduction assay and measuring of the resulting OD490 values before normalizing final values to non-biofilm-forming planktonic cells. Data points represent the mean and the standard error of the mean (n = 72). Differences between groups were tested for significance using Dunnett’s multiple comparison test. ****, p-value <0.0001. ***, p-value <0.001. **(c)** A *C. albicans* CRISPRa strain overexpressing *ALS1* ~6-fold and a non-targeting CRISPRa strain were grown in human urine supplemented with fibrinogen and BSA at 37°C for 48h to simulate a catheterized bladder environment. The robustness of biofilms formed by both strains was measured by reading the OD595 value following a crystal violet assay. Data points are combined from three independent experiments where n = 32 for the non-targeting CRISPRa control and n = 48 for the CRISPRa *ALS1* overexpression strain. Differences between groups were tested for significance using the Mann-Whitney U test. ****, p-value <0.0001. **(d)** Global gene expression of a CRISPRa strain overexpressing *ALS1* ~25-fold, as well as a non-targeting CRISPRa control strain, was measured using RNA-Seq. Differential gene expression analysis showed that *ALS1* had a log2-fold change of ~3.3 in the *ALS1* overexpression strain compared to the CRISPRa control strain, with an adjusted p-value of 3.07E-142, and showed no additional transcripts that had differential expression with an adjusted p-value of less than 0.03.

To confirm phenotypes associated with overexpression of *ALS1*, we performed two distinct biofilm growth assays that compared the *ALS1* CRISPRa strain to the CRISPRa control strain. In the first assay, we allowed biofilms to establish over 24h in flat-bottom 96-well polystyrene plates, removed planktonic cells, and quantified the metabolic activity of the biofilm via XTT reduction, relative to the planktonic cell population. We found that the *ALS1* CRISPRa strain formed significantly more robust biofilms, relative to the control strain (*P* < 0.0005; **FIGURE 4b**). Interestingly, biofilm growth was not correlated with *ALS1* overexpression levels, as the strain with ~25-fold *ALS1* overexpression formed less robust biofilms compared with the strain expressing ~6-fold more *ALS1*. This suggests that very high levels of *ALS1* may ultimately impair biofilm formation, or that the level of *ALS1* expression varies in the conditions used for RT-qPCR analysis compared with biofilm growth, perhaps due to condition-specific repositioning in the TSS.

Next, to mimic conditions associated with fungal biofilm formation in catheter-associated urinary tract infections (CAUTIs), we used a urine-based assay and analyzed biofilm formation of a *C. albicans* strain overexpressing *ALS1*, relative to its control strain, using crystal violet staining. For this assay, we followed up on the *ALS1* CRISPRa strain with ~6-fold overexpression of *ALS1*, which had more robust biofilm growth than the control strain (**FIGURE 4a,b**). The *C. albicans* strains were incubated for 48h in human urine supplemented with bovine serum albumin (BSA) in fibrinogen-coated 96-well plates. Fibrinogen has been shown to be a critical protein for pathogen binding in CAUTIs, by providing a scaffold for biofilm formation and a nutritional source for pathogens (87–89). The *ALS1* CRISPRa overexpression strain showed significantly greater biofilm formation compared to the corresponding wild-type strain (*P* < 0.0001; **FIGURE 4c**). This difference in biofilm formation suggests that *ALS1* promotes binding to fibrinogen and plays a vital role in biofilm formation in urine.

To assess the specificity of this CRISPRa system and ensure no off-target effects from the dCas9-sgRNA complex, we monitored global changes in transcription in the *ALS1* CRISPRa strain via RNA-sequencing (RNA-seq). Since *ALS1* is a member of the large agglutinin-like sequence (ALS) gene family (90, 91), it may be a likely candidate for any potential off-target effects. We also considered *ALS1* as a strong candidate for this RNA-seq off-target profiling since *ALS1* is not known to be a transcriptional regulator, and therefore its overexpression is unlikely to lead to direct downstream transcriptional alterations that would obscure our analysis of CRISPRa off-target effects. Therefore, we performed RNA-seq on the validated CRISPRa strain overexpressing *ALS1* by ~25-fold as well as on a control strain containing the dCas9 cassette and a non-targeting sgRNA when grown in standard laboratory conditions. This analysis confirmed the specific overexpression of *ALS1* (log2 fold change ~3.3 via RNA-seq analysis, adjusted *P*-value 3.07E-142; **FIGURE 4d, TABLE S2**). Five additional transcripts were identified as significantly modulated (adjusted *P*-value < 0.05, absolute log2 fold change > 1; **TABLE S2**). However, their altered expression was much less significant than *ALS1* (**FIGURE 4d**), none of them having an adjusted P-value of less than 0.03, differing from the adjusted P-value of 3e-142 for *ALS1*. We suspect these additional transcripts’ altered expression is likely unrelated to the CRISPRa system based on the fact that 4/5 genes were down-regulated instead of up-regulated, and none of the genes are in proximity to the *ALS1* locus, nor do they appear to share any sgRNA target similarities in their promoter regions. Together this suggests that this CRISPRa system is likely specific with regards to gene targeting.

## Discussion

Here, we present the development, validation, and application of a single-plasmid-based CRISPRa platform for efficient genetic overexpression in *C. albicans*. We demonstrate the ability of this system to overexpress genes ~2-20-fold, and recapitulate important biological phenotypes associated with genetic overexpression, including biofilm formation and antifungal drug resistance. In addition, we assess sgRNA design principles based on predicted TSS, and validate the specificity of this system by monitoring off-target effects via RNA-seq. Together, this work demonstrates an effective, efficient, and specific tool for genetic overexpression in *C. albicans* to enhance the available techniques for genetic modulation in this important fungal pathogen.

Studying genetic overexpression is an important strategy for functional genetic analysis, as overexpressing genes can mimic natural biological states where genes are expressed at high levels. In *C. albicans*, there are numerous scenarios in which the overexpression of genes occurs, including mutations that modify the function of transcriptional regulators and promote the expression of downstream genes (36, 92), as well as aneuploidies such as trisomies (38, 93–98), isochromosome formation (40, 94), and copy number variations (45, 93). These overexpression events are frequently detected amongst *C. albicans* clinical isolates (93, 94, 98) and can play an important role in drug resistance (40, 45, 94, 95), host adaptation (38, 99), and other stress tolerance phenotypes (100, 101). For example, amplification of part or all of chromosome 5, via trisomy or isochromosome formation has been tightly linked to the development of resistance to azole antifungals in *C. albicans* via the increased copy number of genes encoding the azole target, Erg11, and the transcriptional regulator of drug efflux, Tac1 (40). The prevalence and importance of these phenomena associated with increased gene expression highlight the relevance of developing new tools to targetedly assess phenotypes associated with genetic overexpression in *C. albicans*.

Harnessing the cell’s natural transcriptional machinery to increase gene expression with CRISPRa also leads to a unique set of obstacles. Off-target effects as a result of promiscuous dCas9 binding in the cell presents the possibility of complications if the dCas9 is also targetting other regions in the genome. While our RNA-seq results suggest that our CRISPRa system is specific when paired with one sgRNA targeted to the *ALS1* gene, there is still the possibility of off-target effects occurring in different CRISPRa constructs, especially as binding specificity with CRISPR-Cas9 systems is highly dependent on the exact sgRNA sequence in use (102). Using complementary genetic tools to confirm phenotypes observed from CRISPRa strains can help mitigate this. One phenomenon shared among many different CRISPRa activation domains, including VPR, is a reduced capacity to induce overexpression in genes that are already expressed at a high levels (103). Incidentally, the gene that we achieved the lowest overexpression with across all of our testing, *FGR28*, had the lowest basal expression (data not shown) from our RT-qPCR experiments, opposing the idea that genes that are normally expressed at low levels tend to be more easily overexpressed with CRISPRa systems. However, efficiency of the sgRNA molecule and its exact position in the target region of the gene of interest, as well as a host of promoter dynamics, can also influence CRISPR-Cas9 efficacy (69, 104, 105). This illustrates why multiple sgRNAs need to be designed when targeting a gene with CRISPRa to maximize successful overexpression of the target gene. While the rapid and inexpensive nature of CRISPRa strain construction easily justifies the need to design several sgRNA molecules for any given gene, broader limitations as a result of the specific “NGG” PAM recognition site by traditional *Streptococcus pyogenes* Cas9 proteins, like the one used here, leads to the restraint that on rare occasions it may not be possible to produce any highly-efficient sgRNA molecules in a suitable region of the promoter (11). Since this PAM restriction is a feature shared among many CRISPR-Cas9 systems, it may therefore be prudent to further expand the CRISPR toolbox in *C. albicans* by introducing novel Cas9 variants with differing PAM recognition sites into our system, or by constructing novel CRISPR systems exploiting alternative Cas proteins with different and broader PAM recognition capacities, offering the ability to target regions in the genome that cannot be manipulated with current CRISPR-Cas9 tools in *C. albicans* (106, 107).

While large-scale CRISPRa libraries have not been generated in *C. albicans*, their application in other organisms, namely mammalian cell lines, has enabled important discoveries in cancer biology, stem cell reprogramming, cellular fitness, and drug resistance (30, 67, 108–113). The development of CRISPRa systems in *C. albicans* could enable the development of larger overexpression libraries for functional genomic screens, to complement existing fungal gene deletion and depletion libraries (114–117) that have been subject to robust screening and analysis under diverse conditions (118), to reveal critical new insight into fungal biology (119, 120), pathogenesis (115, 120–124), and mechanisms of drug and stress tolerance (125–128).

Currently, non-CRISPR-based overexpression libraries have been constructed and screened in *C. albicans*, revealing novel insights into gene function related to hyphal morphogenesis (60), cell cycle progression (129), drug tolerance (130), and biofilm formation (131). Previous comparisons done between CRISPRa and ORF-based overexpression libraries such as these in other species have highlighted the potential for these two complementary strategies to both confirm the results from one another as well as unveil significant genetic hits distinct from each other, even when libraries share similar gene targets (67). Compared with other overexpression methods, the ability of the CRISPRa system to target any specified region in the genome also confers the unique and powerful ability of allele-specific targeting which could potentially be used to study heterozygosity in *C. albicans*, an aspect of its genomic makeup that is critical to its pathogenicity (132). CRISPRa also offers the advantage of being easily scalable into a pooled format whereby hundreds or thousands of unique plasmids and resulting mutant strains can be constructed simultaneously, and mutant libraries in different strain backgrounds can therefore be quickly constructed (133, 134). Non-CRISPR overexpression libraries that have previously been assembled in *C. albicans* have the potential weakness of missing important phenotypes during screening if the level of overexpression achieved in each mutant strain is too low (46). The ability of our system to cause upwards of 25-fold overexpression in gene targets is thus an important feature. Furthermore, corresponding phenotypes in mutant strains may only be observable if the level of overexpression closely mimics the level of upregulation of the gene induced by the cell in other relevant stress conditions naturally (46). Future iterations of this CRISPRa system could also exploit an inducible dCas9-VPR construct for tunability of gene expression. It is therefore imperative to employ tools such as CRISPRa that possess the advantage of being inherently flexible and easily tunable in order to study genetic overexpression thoroughly.

The application of similar CRISPRa systems to other fungal organisms would lend further insight into fungal biology across this kingdom. Many CRISPR tools have been developed in a diversity of fungal organisms (17, 22, 135–139), including numerous non-*albicans Candida* species (11, 140–146), and other prominent pathogens such as *Cryptococcus neoformans* (147–149) and *Aspergillus fumigatus* (150–153). These CRISPR platforms could be adapted to similar CRISPRa tools in these organisms, with many possible applications for the analysis of pathogen biology or drug target identification (49–52, 154).

Genetic overexpression systems also have many useful applications for industrially-important fungi, as targeted genetic overexpression can be exploited for metabolic engineering to enhance the production of commercially-important metabolites (136, 155–157). CRISPR techniques have already been adapted to numerous industrially-relevant fungi (157–160), and CRISPRa has recently been optimized for use in the filamentous fungi *Aspergillus nidulans* and *Penicillium rubens* (161, 162). These CRISPRa systems were applied to activate transcriptionally-silent biosynthetic gene clusters in these filamentous fungi, with important applications for the discovery of novel bioactive molecules (161, 162). As many *Candida* species play important industrial roles in manufacturing metabolites for food and pharmaceutical industries, as well as remediating wastewater via hydrocarbon degradation (163–167), species-specific adaptations to the CRISPRa tool presented here could produce optimized *Candida* strains for valuable biomanufacturing and bioremediation applications.

## Materials and Methods

### Plasmid design and cloning

The plasmid backbone used in this study was the *C. albicans*-optimized CRISPR-dCas9 plasmid (pRS143, Addgene #122377) used in our previous study (26), containing the *NEUT5L* integration site, sgRNA cloning site (*SNR52* promoter, SapI cloning locus, and sgRNA tail), and *dCAS9*. The dCas9-VPR fusion construct was generated via Gibson assembly. The VPR tripartite complex was codon-optimized for *C. albicans* expression and synthesized as gBlocks gene fragments from Integrated DNA Technologies (IDT). This gene fragment was cloned with Gibson assembly into the *dCAS9* plasmid backbone. We have made the CRISPRa (dCas9-VPR) plasmid available via Addgene (reference #182707). All plasmids are listed in **TABLE S1.**

### sgRNA design and cloning

The sgRNA CRISPR RNAs (crRNA; 20 nucleotide sequence complementary to the target DNA genomic DNA) were designed based on efficiency and predicted specificity via the sgRNA design tool Eukaryotic Pathogen CRISPR gRNA Design Tool (EuPaGDT; http://grna.ctegd.uga.edu) (168). These sgRNA crRNA N20 sequences were cloned into the dCas9-VPR CRISPRa plasmid at the sgRNA cloning locus (SapI cloning site, flanked by *SNR52* promoter, and sgRNA tail) using Golden Gate cloning (169), as previously described (35, 170). crRNA N20 sequences were obtained as two oligonucleotides from IDT in forward and reverse complement orientation, each containing a SapI cloning site, and were reconstituted to 100μM in a nuclease-free duplex buffer from IDT. Equal volumes of the two complementary oligonucleotides were combined after being heated separately at 94°C, and then duplexed by heating to 94°C for 2 min and cooling to room temperature. The duplexed fragment was then cloned into the CRISPRa plasmid with the following Golden Gate cloning reaction: 10μl miniprepped CRISPRa plasmid, 1μl duplexed oligonucleotide, 2μl 10X CutSmart buffer, 2μl ATP, 1μl SapI, 1μl T4 DNA ligase, and 3μl nuclease-free water. This mixture was incubated in a thermocycler under the following cycling conditions: (37°C, 2 min; 16°C, 5 min) for 99 cycles; 65°C, 15 min; 80°C, 15 min. After cycling, 1μl of additional SapI was added to each reaction mixture, and the reaction was incubated at 37°C for 1h. All sgRNAs are listed in **TABLE S1.**

### Media and growth conditions

*Escherichia coli* DH5α cells were grown at 37°C in Lysogeny Broth (LB) and LB plates supplemented with 100mg/mL ampicillin (AMP) and 250mg/mL nourseothricin (NAT) for plasmid selection. *C. albicans* cells were grown at 30°C or 37°C in Yeast Peptone Dextrose (YPD) broth and YPD plates supplemented with 250mg/mL nourseothricin (NAT) for plasmid selection.

### Bacterial transformation

High Efficiency 5-alpha Competent *E. coli* (NEB) cells were thawed on ice. 1μl of Golden Gate cloning reagents were added to 50μl of competent cells and incubated on ice for 30 min, heat shocked for 30s at 42°C, and incubated on ice for an additional 5 min. 950μl of SOC outgrowth media was added to each cell culture and incubated at 30°C for 1.5h at 250 RPM. Transformed cells were then plated on LB media containing AMP and NAT and grown at 30°C for 1 day.

### C. albicans transformation

Plasmids were transformed into *C. albicans* via a chemical transformation strategy, as previously described (35, 170). Briefly, miniprepped CRISPRa plasmids were linearized via a 1.5X restriction digest mix using the PacI enzyme. A transformation master mix was prepared as follows: 800μl of 50% polyethylene glycol (PEG), 100μl of 10X Tris-EDTA (TE) buffer solution, 100μl of 1M lithium acetate (LiAc), 40μl of 10 mg/mL salmon sperm DNA, and 20μl of 1M dithiothreitol (DTT). The transformation mix was added to *C. albicans* cells and linearized CRISPRa plasmid and left to incubate at 30°C for 1h, then heat-shocked at 42°C for 45 min. Cells were washed with fresh YPD, then grown in YPD for 4h to allow for expression of the NAT resistance construct. Transformed cells were then plated on YPD media containing NAT, and grown at 30°C for 2 days. All *C. albicans* strains are listed in **TABLE S1.**

### Growth curves

Overnight cultures of *C. albicans* grown in YPD were diluted to an OD600 of 0.05 in fresh YPD and added to a 96-well flat-bottomed plate with a total volume of 200μl per well. Growth was measured at 30°C via optical density at 600nm at 15 min intervals over the course of 24h using an Infinite 200 PRO microplate reader (Tecan), and plates were shaken orbitally for 900s at a 4mm amplitude in between growth measurements.

### Minimum inhibitory concentration (MIC) assays

Fluconazole MIC assays were performed in 96-well flat-bottomed plates. A suspension of 40μg/mL fluconazole was prepared in water, of which 100μl was added to the first column of each plate containing 100μl of culture, to obtain a total volume of 200μl per well. The first column was serially diluted 2-fold across the plate in water. The gradient of fluconazole, therefore, ranged from 20μg/mL to 0μg/mL. Overnight cultures of *C. albicans* grown in YPD were diluted to an OD600 of 0.1 in 2X RPMI 1640 with 40g/L of added D-glucose and mixed into the plates in an equal volume such that the starting OD600 values of each strain were 0.05 and the growth media was diluted to RPMI 1640 with 20g/L of added D-glucose. Plates were incubated at 37°C at 900 RPM, and absorbance values at 600nm were read after 24h using an Infinite 200 PRO microplate reader (Tecan). The amphotericin B MIC assays were also performed in 96-well flat-bottomed plates similar to the protocol described above with a few modifications. A starting concentration of 5μg/mL of amphotericin B was used to create a gradient ranging from 2.5μg/mL to 0μg/mL. Strains were instead grown in YPD media using a starting OD600 of 0.001. Plates were incubated at 37°C statically and absorbance values at 600nm were read after 72h using an Infinite 200 PRO microplate reader (Tecan).

### Biofilm assays

RPMI-based biofilm assays were performed as previously described (171), with minor modifications. Overnight cultures of *C. albicans* grown in YPD were diluted to an OD600 of 0.001 in 5mL of RPMI 1640 with 20g/L of supplemented D-glucose. 100μl of RPMI 1640 was mixed with 100μl of each diluted overnight culture into a 96-well flat-bottomed plate (12 wells per strain). Plates were grown statically at 37°C for 24h to allow biofilms to form. 120μl of planktonic cell supernatant was removed from each well and moved to a new 96-well flat-bottomed plate. Density of the planktonic cells was measured at 600nm using an Infinite 200 PRO microplate reader (Tecan). Original plates from which planktonic cells were removed were washed by adding 200μl 1X PBS to each well with an electronic multichannel pipette at a moderate dispense speed. 1X PBS was then discarded, and this wash step was repeated.

Plates were then left to dry for 1 - 2h. Once biofilms were dry, 90μl of 1mg/mL XTT (prepared in 1X PBS and centrifuged to remove sediment prior to use) and 10μl of 0.32mg/mL PMS (prepared in water) were added to each well. Plates were incubated statically at 30°C for 2h to allow biofilms to reduce XTT, measured at 490nm using an Infinite 200 PRO microplate reader (Tecan), and normalized to the growth of planktonic cells harvested previously.

For urine-based biofilm assays, biofilm formations were performed in 96-well flat-bottomed plates. Plates were coated with 150μg/mL of fibrinogen and incubated overnight at 4°C. The *C. albicans* strains were cultured at 37°C with aeration in 5mL of YPD broth. The inoculum was normalized to ~1×10^6^ CFU/ml and then diluted (1:10) in human urine (female, pH ~6.5). Urine was supplemented with 20mg/mL of BSA for carbon and nitrogen sources mimicking the catheterized bladder environment. 200μl of the urine with inoculum was incubated in each well of the 96-well plate at 37°C for 48h while static. Following the 48h incubation, the cultures were removed and the wells were washed three times with 200μl 1X PBS to remove unbound fungi. Plates were incubated with 200μl of 0.5% crystal violet for 15 min. Crystal violet was removed, and plates were washed with water and dried. 200μl of 33% acetic acid was added to the wells and 100μl was transferred to a new 96-well plate. Absorbance was measured at 595nm using a SpectraMax ABS Plus microplate reader (Molecular Devices).

### RNA extraction and real-time quantitative PCR

To detect differences in gene expression, overnight cultures of *C. albicans* grown in YPD were diluted to an OD600 of 0.05 in fresh YPD and grown to an OD600 of >0.2 at 30°C. Cultures were pelleted and frozen at −80°C before RNA was extracted. RNA extractions were performed using either a Presto Mini RNA Yeast Kit from FroggaBio (cat. RBYD050) or an RNeasy Mini Kit from Qiagen (cat. 74104). To assess RNA integrity, an RNA ScreenTape assay was performed on all samples using the TapeStation 4150 system following manufacturer’s instructions (Agilent Technologies). Only samples with RIN values of >5.0 were used for RT-qPCR. Synthesis of cDNA from 1000ng (10μl) RNA was performed using a High Capacity cDNA Reverse Transcription Kit from Applied BioSystems (cat. 4368814). Briefly, 20μl reaction mixtures were prepared with 2μl 10X reverse transcription buffer, 0.8μl of 25X dNTPs at a concentration of 100mM, 2μl of random primers, 1μl of MultiScribe Reverse Transcriptase at 50 U/μl, 4.2μl of nuclease-free water, and 10μl of RNA (1μg). Reverse transcription run conditions were as follows: 10 min at 25°C, 120 min at 37°C, 5 min at 85°C. Real-time PCR assays were conducted using a QuantStudio 7 Pro Real Time PCR system from Thermo Fisher Scientific Inc. 20μl reaction mixtures containing 10μl 2X SsoAdvanced Universal Inhibitor-Tolerant SYBR supermix (Bio-Rad, cat: 172-5017), 0.8μl of PCR forward and reverse primer mix at 5μM (final concentration of primer at 200 nM), 4.2μl of water and 5μl of diluted cDNA. The run conditions were as follows: 3 min at 98°C polymerase activation step, followed by 40 cycles of a two-step qPCR (10s of 98°C denaturation, 30s of 60°C combined annealing/extension). Primers are listed in **TABLE S1**. Expression profiling calculations were performed according to the comparative C_T_ method (172). Briefly, expression values for the experimental gene of interest were compared to the housekeeping gene *ACT1* to obtain a ΔC_T_ value within each strain. The ΔC_T_ values in the experimental CRISPRa strains were then compared to the non-targeting CRISPRa control strain to obtain a ΔΔC_T_ value and finally a fold-difference in expression of the experimental gene.

### RNA sequencing

RNA preparation and sequencing were performed as previously described, with minor modifications (171). Strains were grown in YPD at 30°C overnight preceding RNA extraction. RNA extraction, sample QC, library preparations, and sequencing reactions were conducted at GENEWIZ, LLC./Azenta US, Inc. as follows: Total RNA was extracted using Qiagen RNeasy Plus Universal kit following manufacturer’s instructions (Qiagen). RNA samples were quantified using Qubit 2.0 Fluorometer (ThermoFisher Scientific) and RNA integrity was checked with 4200 TapeStation (Agilent Technologies). ERCC RNA Spike-In Mix kit (cat. 4456740) from ThermoFisher Scientific, was added to normalized total RNA prior to library preparation following the manufacturer’s protocol. The RNA sequencing library was prepared using the NEBNext Ultra II RNA Library Prep Kit for Illumina using the manufacturer’s instructions (New England Biolabs). Briefly, mRNAs were initially enriched with Oligod(T) beads. Enriched mRNAs were fragmented for 15 min at 94 °C. First-strand and second-strand cDNA were subsequently synthesized. cDNA fragments were end-repaired and adenylated at 3’ends, and universal adapters were ligated to cDNA fragments, followed by index addition and library enrichment by PCR with limited cycles. The sequencing library was validated on the Agilent TapeStation (Agilent Technologies), and quantified by using Qubit 2.0 Fluorometer (ThermoFisher Scientific) as well as by quantitative PCR (KAPA Biosystems). The sequencing libraries were multiplexed and clustered onto a flowcell. After clustering, the flowcell was loaded onto the Illumina HiSeq 4000 or equivalent instrument according to the manufacturer’s instructions. The samples were sequenced using a 2×150bp Paired End (PE) configuration. Image analysis and base calling were conducted by the Illumina Control Software.

Raw sequence data (.bcl files) generated from the Illumina instrument was converted into fastq files and de-multiplexed using Illumina bcl2fastq 2.20 software. One mis-match was allowed for index sequence identification. Sequence reads were trimmed to remove possible adapter sequences and nucleotides with poor quality using Trimmomatic v.0.36, and trimmed reads were mapped to the candida_albicans reference genome available on ENSEMBL using the STAR aligner v.2.5.2b. The STAR aligner is a splice aligner that detects splice junctions and incorporates them to help align the entire read sequences. BAM files were generated as a result of this step. Unique gene hit counts were then calculated by using featureCounts from the Subread package v.1.5.2. The hit counts were summarized and reported using the gene_id feature in the annotation file. Only unique reads that fell within exon regions were counted. After extraction of gene hit counts, the gene hit counts table was used for downstream differential expression analysis. Using DESeq2, a comparison of gene expression between the control and experimental groups of samples was performed. The Wald test was then used to generate p-values and log2 fold changes. Genes with an adjusted p-value < 0.05 and absolute log2 fold change > 1 were called as differentially expressed genes for each comparison. The data have been deposited in NCBI’s Gene Expression Omnibus (GEO) (Edgar *et al*. 2002) and are accessible through GEO Series accession number GSE201904 (https://www.ncbi.nlm.nih.gov/geo/query/acc.cgi?acc=GSE201904).

### Data availability

The *C. albicans* CRISPRa plasmid backbone was deposited to Addgene as plasmid number #182707. RNA sequencing data was deposited in NCBI GEO repository (accession number GSE201904; https://www.ncbi.nlm.nih.gov/geo/query/acc.cgi?acc=GSE201904).

## Acknowledgments

We thank Shapiro lab members, past and present, for helpful discussions and support of this project, as well as Dr. Anna Selmecki and Dr. Petra Vande Zande (University of Minnesota) for helpful feedback. We thank Dr. Zhenguo Lin (Saint Louis University) for helpful discussions on TSS prediction, and Kieran Shah for programming support on this project. We thank Jing Zhang from the University of Guelph Advanced Analysis Centre for RT-qPCR support. This work was supported by an NSERC Discovery Grant (RGPIN-2018-4914), an Ontario Early Research Award, and a CIFAR Azrieli Global Scholar award to RSS. N.C.G, L.W, and M.F. are EvoFunPath Fellows (NSERC CREATE), L.W. is supported by an Ontario Graduate Scholarship, M.F. is supported by an NSERC-CGS-M scholarship, and R.O.C.L was supported by a MITACS Globalink Research Internship. A.C. is supported by a Career Award for Medical Scientist from the Burroughs Wellcome Fund. Work by A.L.F.M. and F.H.S.T. was supported by institutional funds from the University of Notre Dame as well as National Institute of Health grant R01-DK128805 (to A.L.F.M. and A.A.L.), and the Arthur J. Schmitt Presidential Leadership Foundation (to A.L.L).

## Notes

### Competing Interest Statement

The authors have declared no competing interest.

### Summary of Updates

This version of the manuscript has revised Figure 3

